# The COMPARE Data Hubs

**DOI:** 10.1101/555938

**Authors:** Clara Amid, Nima Pakseresht, Nicole Silvester, Suran Jayathilaka, Ole Lund, Lukasz D. Dynovski, Bálint Á. Pataki, Dávid Visontai, Matthew Cotten, Basil Britto Xavier, Blaise Alako, Ariane Belka, Jose J. L. Cisneros, George B. Haringhuizen, Peter W. Harrison, Dirk Höper, Sam Holt, Camilla Hundahl, Abdulrahman Hussein, Rolf S. Kaas, Surbhi Malhotra-Kumar, Rasko Leinonen, David F. Nieuwenhuijse, Nadim Rahman, Carolina dos S Ribeiro, Jeffrey E. Skiby, József Stéger, János M. Szalai-Gindl, Martin C. F. Thomsen, István Csabai, Marion Koopmans, Frank Aarestrup, Guy Cochrane

## Abstract

Data sharing enables research communities to exchange findings and build upon the knowledge that arises from their discoveries. Areas of public and animal health as well as food safety would benefit from rapid data sharing when it comes to emergencies. However, ethical, regulatory, and institutional challenges, as well as lack of suitable platforms which provide an infrastructure for data sharing in structured formats often lead to data not being shared, or at most shared in form of supplementary materials in journal publications. Here, we describe an informatics platform that includes workflows for structured data storage, managing and pre-publication sharing of pathogen sequencing data and its analysis interpretations with relevant stakeholders.

## Introduction

Whole genome sequencing (WGS) methods show increasing appeal for those involved in pathogen diagnosis, tracking and research. This includes institutions responsible for public or animal health and food safety, clinical microbiology laboratories delivering patient diagnostics, treatment advice and hospital epidemiology support, as well as industries working to avoid disease outbreaks or those developing novel solutions for detection or treatment. Last but not least, this field is of interest to the general scientific community engaged in the mechanistic exploration of pathogen biology and epidemiology. Indeed, advances in DNA sequencing technologies and the lowering of the associated costs, have led to an increasing uptake of whole genome sequencing (WGS) and next generation sequencing (NGS) in these areas. However, NGS methods are data-intensive with complex analytical requirements, such that their adoption places demands on local, national and global infrastructure for the management, sharing and effective exploitation of NGS-derived pathogen data. This particularly applies when it comes to supporting outbreak investigations, where time is of essence, and pathogen genome characterisation can lend support to public health officials charged with outbreak management. As pathogens do not respect (national or institutional) borders, the need for rapid sharing of data is increasingly endorsed (1). The World Health Organisation for example has repeatedly advocated open sharing of pathogen genetic sequences as well as the knowledge and benefits resulting from the genetic data (https://www.who.int/blueprint/meetings-events/meeting-report-pathogen-genetic-sequence-data-sharing.pdf). However, data sharing has faced some resistance as capacities and capabilities to generate the data differ greatly between countries and regions. In addition, political, ethical, administrative, regulatory and legal barriers can cause major delays in the sharing of data, making it almost practically impossible (2–3).

The European Union’s HORIZON 2020 initiative COMPARE (http://www.compare-europe.eu/) set out in 2015 to rise to the challenges brought by the advent of NGS methods in pathogen surveillance and research (3). COMPARE aims to provide solutions for barriers to implementation of NGS spanning all domains (clinical, animal health and food) and pathogens (viruses, bacteria and eukaryotic parasites), and includes low resource settings. As a guiding principle, COMPARE aimed to setup a user-friendly data upload, analysis and sharing platform, where all NGS data with relevant minimum metadata are eventually shared in the public domain through the permanent repositories of the International Nucleotide Sequence Database Collaboration (INSDC, 4, http://www.insdc.org/). This model addresses the following key barriers: 1) long term data storage of high density data; 2) bioinformatic analysis of data for users without in house expertise; 3) a trusted environment for prepublication sharing of data with relevant stakeholders; 4) data structuring for searchability and re-useability by other scientists and visualisation of data. For this, COMPARE has developed and tested a number of informatics platforms and bioinformatic workflows by which users can manage, share, analyse and interpret NGS pathogen data.

In this paper, we introduce the COMPARE Data Hubs (CDHs), designed to be used as focal points for collaborative data sharing and analysis around which user communities engage, and the centrepiece of COMPARE’s informatics platform. Our discussion covers overall design of the system, highlights our principles of standards-compliance and open data, and details the anatomy of a CDH. We provide examples of CDHs in use and describe how users can access the system. Finally, we lay out future plans for further developments and applications of the CDH system.

## COMPARE Data Hub design

The CDHs offer a complete platform through which a group of collaborating users can share, analyse and interpret their data. A typical use of the system starts with a data provider reporting sequence read data and choosing the appropriate CDH for sharing (see figure I). Defined autonomous processes detect these incoming datasets and route them through an appropriate analysis workflow, as selected when the CDH was configured. The results of the analysis, themselves, further data (albeit derived), automatically flow back into the CDH when workflow compute processes reach completion. Finally, users with access to the CDH can then search for data in the CDH, explore and visualise analysis results and select data sets for retrieval.

**Figure I:**
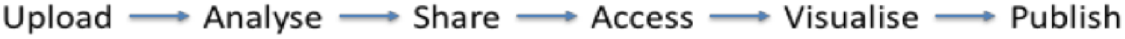
Example CDH user workflow

Following our design principles (see Box I), the CDHs have been implemented as a modular system that centres around a data storage module with connected cloud compute (see figure II). Surrounding these components are a number of tools and interfaces that support steps in the user workflow, such as those described above. These tools and interfaces are brought together within the Pathogens Portal (PP) website to provide a single point of entry for users making use of the CDHs (https://www.ebi.ac.uk/ena/pathogens). Data reporting tools are offered that support upload and validation of data and metadata; these span the spectrum from an intuitive web “drag and drop” system for bacterial isolates to Application Programming Interfaces (APIs) that allow laboratories to connect data reporting directly to their existing informatics systems. The control of data sharing by data providers is supported directly within the PP (under the “Share” tab) or through an API (https://www.ebi.ac.uk/ena/portal/ams/). Data search functions are also presented directly in the PP (under the “Search” tab) and through an API (https://www.ebi.ac.uk/ena/portal/api/). Exploration and visualisation are available in the PP under the “Explore” tab and data download tools are available variously within the PP/API and through a number of download management tools for public data. The latter include the ENA browser and services providing high-volume data access such as the ENA File Downloader (https://github.com/enasequence/ena-ftp-downloader).

**Figure II:**
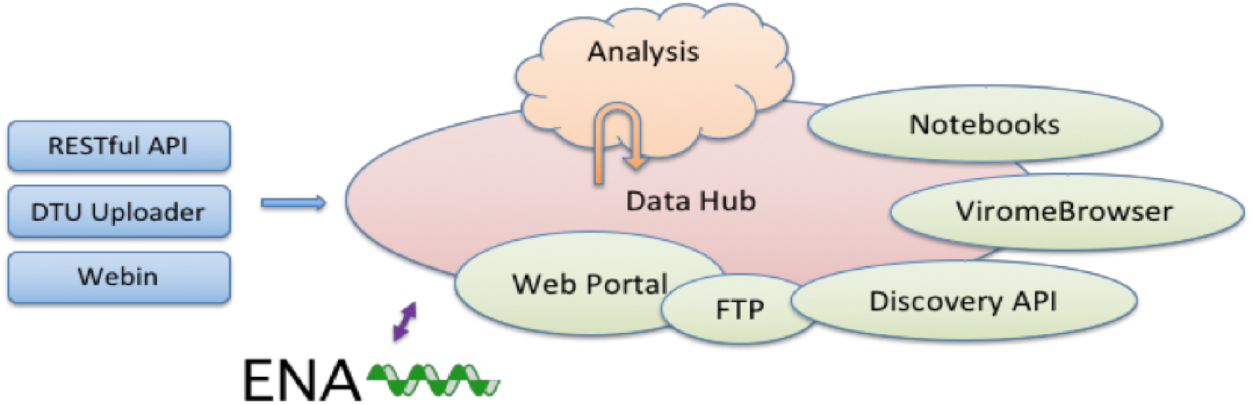
System architecture

CDHs are modular and flexible; no two CDHs are configured or used in exactly the same way. At time of set up of a given CDH, as a user group comes together and agrees on the work that they will carry out, a new CDH is configured with all of the appropriate components that are required for the CDH. Component choices include metadata checklists (that will be used to validate incoming data sets and define searchable fields), choice of analysis workflow and selection of appropriate visualisations. Other component choices (such as data reporting interfaces) can be made at the time of use, or based on individual (user) preference to allow, for example, one data provider to a hub to use API-based data reporting and another to use a web interface.

In designing the CDH system, we have paid particular attention to the reuse and adaptation of existing open data infrastructure where possible. The most notable example of this is the reuse of the European Nucleotide Archive (ENA, 5, https://www.ebi.ac.uk/ena) and many of its surrounding tools for COMPARE’s integration, storage, presentation and retrieval of data. As the European node in the long-established global public databases of record for sequence data, the INSDC (4), the ENA maintains the comprehensive public record for pathogen sequence data. A number of criteria led us to build around ENA: we needed to be as cost-effective as possible in our engineering by leveraging past investment; we sought the high sustainability, particularly for long-term data preservation, that ENA offers; there is a frequent user need to be able to operate on their own data in the context of all available public data; and use of the ENA system at the heart of the CDHs, along with a commitment to structured data sharing (see Box I), assures that data and metadata are appropriately compliant with publication requirements for when full publication of the data is required.

### Box I

#### Design principles

- ***Responsible open data:*** COMPARE strongly advocates open science and promotes open scientific data; a true understanding of the pathogens in global circulation and an ability to model and predict can only be possible through a comprehensive view across all previously observed pathogens; for NGS-based methods, what is observed today in surveillance and diagnostics becomes the reference point for tomorrow’s outbreak investigation; all data shared through CDHs are thus ultimately destined for public view.
- ***Structured data sharing:*** in order to aid discovery, interoperability and to drive at maximum value, data sharing in the CDHs has structure and formality; data are validated at the time of reporting against defined and published specifications; similarly, metadata associated with incoming data sets are validated against the appropriate standards; new data are thus integrated into the system in structured form to support fully searchability, interoperability and readiness for computational analysis processes.
- ***Data standards:*** international community standards result from expert engagement in defining appropriate structures and elements of metadata that are required for integrity, utility and impact of shared data; CDHs build in these standards, in particular those of the Global Microbial Identifier initiative (http://www.globalmicrobialidentifier.org/) and the MIxS family of standards (6) from the Genomics Standards Consortium (https://press3.mcs.anl.gov/gensc/); delivered through a checklist and data file validation system, standards compliance is made practical for data providers and impactful for data consumers.
- ***Configurability:*** just as no two collaborations around pathogen data sharing and analysis are identical, the system must be configurable to allow flexibility and choice for each CDH as to tools for reporting, standards to be followed, analysis workflows to be followed; the set up of a new CDH involves detailed configuration with many possibilities for data providers and consumers.
- ***Neutrality:*** trust in the neutrality of the system into which the world places its pathogen data is critical; providing the CDHs and underlying data storage from an international organisation, the European Molecular Biology Laboratory (EMBL, https://www.embl.de/aboutus/general_information/), with more than 20 member states, assures this neutrality.
- ***Extensibility:*** WGS methods are in rapid development from sampling approaches, through sequencing technologies to computational analyses; the CDHs are built with extensibility by design, giving the ability and flexibility to support new sequencing platforms, new sequencing approaches, the latest analysis workflows.
- ***Systematic autonomous analysis:*** high-throughput WGS analysis requires minimal human intervention to minimise time to results and to deliver comparable and interoperable outputs; in the COMPARE system, human interaction is focused into the CDH setup and configuration phase, allowing full autonomy when production operations begin.
- ***Discovery and navigation options:*** search across shared data and analysis results are central to users’ interpretation of pathogen WGS data sets; the CDHs offer powerful web and programmatic metadata search and navigation options; ongoing work to extend and improve search includes deeper indexing of analysis results and addition of sequence similarity search.
- ***Agile visualisations:*** interpretation of analyses requires iterative exploration of data and refinement of visualisation; using the Jupyter Notebook system (https://jupyter.org/), the CDHs support agile addition and modification of visualisations as needs of the CDH’s users emerge.
- ***Sustainability:*** a platform for pathogen WGS data sharing and analysis is needed for the long-term and beyond any one project; building on the established and sustained bioinformatics infrastructure, in particular the ENA, preservation of CDH data is assured and key services around the CDH have the greatest possible future-proofing.
- ***Comprehensive:*** the diversity of WGS-based pathogen methods to sequencing platforms, use of these platforms and computational analyses must be served; the CDHs support data sharing across all major current sequencing platforms and library methods and leverage ongoing investment in engineering around emerging platforms; our cloud-based approach to analysis, in which new computational workflows can be installed allows future computational approaches to be implemented as these emerge.
- ***Interfaces:*** just as users differ in the scale of their data operations and the level of expertise in informatics, platforms must offer a variety of appropriate interfaces; supporting all CDH functions through both web and programmatic interfaces, our systems supports the spectrum of users.
- ***Support:*** users require guidance, training and direct human support; the CDHs are documented with user guides and training materials; in-person training is offered and we operate an expert email Help desk around the CDHs.

## Anatomy of a Data Hub

### Upload and Standards

The COMPARE platform offers a number of data reporting tools that support upload and validation of sequencing and metadata into the CDHs.

One of the prerequisites to structured data sharing is that metadata accompanying uploaded data must fulfil an appropriate standard (see Box I). In the case of samples the choice of standard depends on the sample provenance, i.e. the type of sample being described. The richer the sample metadata, the higher the downstream usability of associated data. The metadata standards are represented as checklists in ENA that allow various types of validations (e.g. regular expressions, use of controlled vocabulary, ontologies). Examples of metadata checklists developed around pathogens and currently available for use within CDHs are:

Prokaryotic pathogens: http://www.ebi.ac.uk/ena/data/view/ERC000029

Influenza virus : http://www.ebi.ac.uk/ena/data/view/ERC000032

Virus pathogen: http://www.ebi.ac.uk/ena/data/view/ERC000033

Sewage: http://www.ebi.ac.uk/ena/data/view/ERC000036

Parasites: https://www.ebi.ac.uk/ena/data/view/ERC000039

### DTU Uploader

The DTU Uploader is one of the web-based data reporting tools offered in the COMPARE platform. The Uploader was created to facilitate submissions of sequencing data of various formats (e.g. FASTQ, FASTA) by scientists working in [remote] wet-laboratories and hospitals. The main goals behind the design of the Uploader were providing users with an easy upload tool for large data files and metadata describing those files, validating content/metadata of the files before and after submission, providing resumable uploads in case of upload failures, as well as sharing uploaded data with ENA -if explicitly specified by user - to allow structured data sharing within the CDHs (see Box I). Moreover, additional validation is executed against the ENA submission API that validates samples to be submitted. In case of failing to comply with ENA validations the user receives an automatic error reporting email in order to fix the issues and resubmit. Uploading takes 3 steps. The first step is to download a metadata *Excel* file from the Uploader interface and filling it out with a minimum set of mandatory information about the upload, such as the sample name and sequenced file(s) associated with the sample. The second step provides a drag and drop functionality for the data and metadata file(s) to be uploaded. The final step is to initiate the upload process, by clicking an upload button.

### Webin interactive

Webin Interactive is ENA’s web-based submission tool (https://www.ebi.ac.uk/ena/submit/sra/#home) available in the COMPARE platform. This tool scales up to large data sets (several hundred of isolates) and allows both completing a form interactively (only recommended for small data sets) and downloading a spreadsheet for off-line completion and re-upload. Data submissions to ENA through Webin Interactive require the registration of a Webin account (Webin-XXXX) and a subsequent four steps that include 1) study and 2) sample registrations, 3) uploading files and 4) submitting uploaded files. Data files can be uploaded using the Webin File Uploader, downloadable from Webin Interactive (https://ena-docs.readthedocs.io/en/latest/upload01.html#using-webin-file-uploader) but also FTP or Aspera protocols (https://ena-docs.readthedocs.io/en/latest/upload_01.html#upload-files). The submission step 4 following data upload allows the user to describe their experiments and the uploaded data files by for example specifying the file format (e.g. FASTQ, BAM, CRAM), sequencing platform or library protocols, and referencing the samples.

### Webin-CLI

The Webin Command Line Interface (Webin-CLI) is a further tool released in late-2018 by ENA for use on both UNIX-based and Windows platforms (https://ena-docs.readthedocs.io/en/latest/cli.html#command-line-submissions). Webin-CLI is provided in the form of a standalone executable JAR file, which can be downloaded from https://github.com/enasequence/webin-cli/releases and run from a UNIX terminal or Windows command prompt. In similar fashion to Webin Interactive or DTU Uploader, here the user must also provide appropriately formatted sequence data files (e.g. FASTQ, BAM). When using Webin-CLI a ‘manifest’ file must be created describing the experimental metadata and naming the sequence files in a simple tab-separated format. The user then constructs a command, including the name of the manifest file and Webin account login credentials. When the command is run, the files are validated and uploaded, after which accession numbers are output.

#### $ webin-cli -context reads -manifest reads_manifest.txt -submit -userName Webin-XXXX -password XXXXX

##### An example of a Webin-CLI command

The primary advantage of Webin-CLI over other submission routes is pre-submission validation: while historically, files have been uploaded and registered before their validity is checked, Webin-CLI validates files before allowing their upload to commence. Hence also, a preceding data file upload to ENA (step 3 in Webin Interactive) is not required here. Any errors are reported directly to the user. From the user perspective this allows greater self-sufficiency, as errors can be fixed at an earlier stage and directly by the submitter without the need to contact the ENA Helpdesk. Contacting helpdesk remains an option if the user can’t solve the problem alone. From the perspective of the database, the benefits include a reduction of Helpdesk overheads and a much-reduced proportion of invalid submissions, making it easier to support ever-growing volumes of submitted data.

### Sharing

Once data providers have uploaded their validated data through one of the described submission routes they then have full control over which of the uploaded studies to share with which CDH. The CDH sharing preferences can be changed by the data provider from within the PP itself (under the “share” tab; https://wwwdev.ebi.ac.uk/ena/pathogens/login?returnUrl=%2Fshare) or programmatically through the authenticated API (https://www.ebi.ac.uk/ena/portal/ams/webin/auth). The datasets are listed by their study accession and title, and the same dataset can be shared across multiple different CDHs if desired. This fine-grained, data provider lead, control was crucial for the rapid distribution of data through the COMPARE platform, particularly as a number of the CDHs have established automated analysis workflows that trigger as soon as new data is shared with the CDH.

### Analysis

A key aim for the COMPARE initiative was to develop and incorporate analytical workflows within a computational workflow engine that would automate the data selection, analysis pipelines and subsequent archival of results for use by the community. To achieve this we developed SELECTA, a rule-based analysis pipeline scheduler that fully automates the operation and automatic submission of generated results to the European Nucleotide Archive for subsequent discovery and retrieval. SELECTA consolidates a number of different analysis pipelines developed within the COMPARE consortium enabling the selected analysis pipeline to be set to automatically run on any newly shared datasets within the CDH. The SELECTA system has fully integrated a virus sequence classification tool, SLIM, the Center for Genomic Epidemiology (CGE) bacterial genome analysis pipeline, BAP, of the Danish technical University, and the University of Antwerp bacterial genome analysis pipeline, UAntwerp Bacpipe. Further pipelines for the comprehensive taxonomic classification of metagenomic sequence reads (RIEMS) are also currently being developed for integration into the SELECTA workflow engine. Each of these workflows is described in more detail below. The output results from each of the pipelines is automatically submitted to the ENA which removes the burden that submitter faces when handling bulk submissions, ensures rapid data processing and consistent data reporting.

#### CGE – bacterial analysis pipeline (BAP)

The CGE workflow is described in Thomsen *et al.* (7). The Docker (https://www.docker.com/; https://opensourceforu.com/2017/02/docker-favourite-devops-world/) based version of the pipeline used in this project uses raw reads from the illumina platform as input; cgMLST and SalmonellaTypeFinder were added to the pipeline (Table I). The pipeline automatically identifies the bacterial species and assembles the genome, if applicable, identifies the multilocus sequence type, plasmids, virulence genes and antimicrobial resistance genes. A short printable report for each sample as a tab-separated file containing all the metadata and a summary of the results for all submitted samples can be downloaded. The source code is available at https://bitbucket.org/genomicepidemiology/cge-tools-docker/src/master/.

**Table 1:** list of the bacterial lineages for which the CGE analysis pipeline has databases

| Database | Included lineages / plasmids |
| --- | --- |
| PlasmidFinder | Enterobacteriaceae, Acinetobacter baumannii, Enterococcus, Streptococcus and Staphylococcus |
| VirulenceFinder | Escherichia coli, Shigella |
| SalmonellaTypeFinder | Salmonella |
| cgMLST | Escherichia coli, Campylobacter, Listeria, Yersinia, Salmonella |
| MLST | has a database for each scheme in the pubMLST database |
| ResFinder | all databases in the ResFinder database |
| pMLST | IncF, IncHI1, IncHI2, IncI1, IncN, IncAC |

#### SLIM - viral identification pipeline

SLIM is a python-based wrapper for two main functions, *de novo* assembly to contigs and contig classification, and their required substeps. After running a generic quality control on the short read data to remove common adapters, short reads and low quality reads, SLIM performs a *de novo* assembly using SPAdes, described by Bankevich *et al.* (8) on the quality controlled short reads using conditions for either Illumina HiSeq paired-end reads or Ion Torrent single reads. Next, SLIM classifies the contigs based on a translation of all six reading frames of the contig and a usearch (Edgar, 9) screen for homologies to viral proteins. The output of this classification is a single tab-separated value (TSV) table listing each contig showing protein homology above 30% to sequences in virus family-based databases. Contigs showing homology above the threshold to a virus family entry are copied into family-specific fasta files with the contig sequence in the standard 5’ to 3’ orientation of the matched virus. A short annotation of the closest identified match (the INSDC accession number and a truncated ID derived from the INSDC entry) is added to the contig’s fasta id. Contigs that failed to return a homology to any entry in any of the databases are gathered as ‘mystery contigs’. The tool provides the usearch alignment file for each virus family to allow users to verify classifications. Summary details are also collected. The output from each analysis run is a TSV summary table with all classification metrics and a compressed directory containing the summary file, the virus family specific fasta and alignment files and the ‘mystery contig’ files. The TSV summary table is a useful document for further exploration of the classification results. Because of the need to limit user-defined and run-specific parameters, generic conditions were chosen that provide a reasonable de novo assembly performance across a broad range of sequence data types. Thus the results serve as a useful starting point for analysis but can usually be improved by tuning condition for run-specific parameters.

#### BACPIPE - bacterial analysis pipeline

Bacpipe (Xavier, B.B. *et al.,* 2019, in preparation) is a collection of open-access bioinformatics tools carefully designed into a logical workflow for the analysis of microorganism whole-genome sequencing and was developed to mitigate the level of bioinformatics experience required for microorganism genome analysis in a clinical setting. This computationally low-resource bioinformatics pipeline enables direct analyses of bacterial whole-genome sequences (raw reads, contigs or scaffolds) obtained from second and third-generation sequencing technologies. Bacpipe covers the full analysis workflow from read quality assessment, to genome assembly, annotation and finally the identification of resistance and virulence genes. The outbreak module (SNPs and patient metadata) can simultaneously analyse many strains to identify evolutionary relationships and transmission routes. Importantly, parallelization of tools in BacPipe considerably reduces the time-to-result. BacPipe is able to simultaneously analyse numerous strains of bacteria to elucidate their evolutionary relationships and derive a microorganism transmission route. BacPipe was initially validated using a Methicillin-Resistant *Staphylococcus aureus* (MRSA) outbreak WGS dataset amongst different datasets from hospital, community and food-borne outbreaks and from transmission studies of important pathogens demonstrating the speed and simplicity of the pipeline that reconstructed the same analyses and conclusions within a few hours. BacPipe consolidates the analysis results into a single worksheet to aid rapid interpretation by clinicians, making it an ideal tool for WGS data analysis and interpretation for routine patient-care in hospitals and for infection monitoring in public health settings.

#### RIEMS - metagenomic analysis pipeline

Metagenomics over recent years has proved to be a powerful tool for the analysis of microbial communities for both clinical diagnostic and scientific purposes. However, a major bottleneck is the extraction of relevant actionable information from these often huge metagenomics datasets. Reliable Information Extraction from Metagenomics Sequence datasets (RIEMS, 10) was developed to address this challenge by accurately assigning each read in a dataset to a taxonomic group. RIEMS analysis proves to be highly accurate when compared to similar metagenomics tools on simulated sequence reads and in 2011 was used to detect the orthobunyavirus sequence in metagenomics reads prompting the discovery of the Schmallenberg virus.

## Discovery

The raw data from providers and the automatically processed analysis results from SELECTA workflows are discoverable through the PP and associated authenticated APIs. The PP acts as a single access point to the wealth of raw and processed analysis data. The PP includes an advanced search query builder (https://www.ebi.ac.uk/ena/pathogens/search) that assists the user in creating powerful searches to identify datasets and analyses of interest. Similarly the Discovery API has a Swagger (https://swagger.io/) interface to assist users with query construction and access to documentation on the usage and status codes of each of the endpoints (https://www.ebi.ac.uk/ena/portal/api/#!/Portal_API/downloadDocUsingGET). The documentation can also be downloaded in PDF. Both the PP advanced search and Discovery API support user authentication so that search is extended to the Data Hubs that the authenticated user has been granted access to, in addition to any public datasets. For complex API queries that may take longer to complete the user can specify an email notification when the search result is ready for download. For both PP web interactive and API interfaces the returned result can be set to include all searchable fields but can also be customised to include a set of fields of interest only, a particular order of the fields and whether to limit the number of results returned. Please also refer to the ‘Usage and Access’ section.

## Retrieval

Once users have identified datasets of interest they can utilise a range of tools to download the data files. The design of the CDH system to utilise existing technology of the European Nucleotide Archive at EMBL-EBI, gives CDH users access to its range of powerful download applications for programmatic users through its Application Programming Interfaces. For non-programmatic users there are a range of graphical interface options, and for pathogens data this is primarily the ENA File Downloader and ENA Browser tools. The ENA File Downloader (https://github.com/enasequence/ena-ftp-downloader) is a stand alone Java based graphical user interface (GUI) application which allows a user to search directly by either an accession or a search query, or alternatively upload a search report generated from within the PP to initiate the download of those sequences. The download interface supports selecting multiple data files at once, download progress indication and automatic verification that files have been successfully downloaded using MD5 checksums. The interface supports downloads using either File Transfer Protocol (FTP) or for less stable connections IBM Aspera transfer software (https://downloads.asperasoft.com/). The ENA Browser tools (https://github.com/enasequence/enaBrowserTools/) are a set of Python based tools that allow command line based (or programmatic) downloads without requiring scripting knowledge. The tools allow downloading all data for a given accession or data of a particular group for a given accession with minimal effort from the user. Once again the tools support both FTP and Aspera based downloads.

## Exploration

### The Notebooks

Data sharing has significantly advanced several disciplines but sharing of the data analysis process in a reproducible way is somewhat lagging behind. Scientific results are traditionally published as human readable articles, but human language lacks the precision and details of computer codes. Therefore, reproducing results even with the data at hand is, if at all possible, often a long tiresome process. The figures in traditional articles without possibility for zooming or subsetting also hide many of the details present in the data and especially multidimensional data sets are hard to represent in passive two dimensional prints. In the late 80s Mathematica’s first Notebook frontend was released. Since then slowly but steadily other languages like R and Python picked up the concept of “reproducible research” (11). In the last couple of years Jupyter Notebooks (12) - capable of handling Python, R and several other languages - became a standard for data analytics and visualisation. The Notebooks integrate the analysis and visualisation process and produce output that can be rendered as rich interactive web pages. Due to the virtualization techniques it was straightforward to integrate these tools into the CDH (Figure II). We use a collaborative cloud-accessible platform, Kooplex (Visontai, D. *et al.,* 2019, in preparation) to perform the final step of data analysis and visualisation. The rendering of Notebooks for presentation through the PP is automated, with the daily generated Notebooks immediately available for distribution. Notebooks can be rendered within the users web browser or downloaded, daily reports are archived so that previous reports can still be accessed. Two types of Notebooks are available: a simpler, static HTML that can give a quick overview about the content of the Data Hub at any given day, and a more complex version the users can use to filter, sort, arrange and visualise the data in their web browser without writing code. In the current version the primary aim of these Notebooks is to lower the initial effort needed to explore the content of a Data Hub but further development may allow even richer data exploration and visualisation. Exploration and visualisation are available in the PP under the “Explore” tab. Please also see the ‘Usage and Access’ section.

### The ViromeBrowser

Where a broad overview of families or genera is usually sufficient in bacterial metagenomics, analysing the virome requires species or sub-species level resolution of sequence annotations to extract useful information. In addition, the detection of a partial sequence of a highly pathogenic virus can require further investigation, while a large quantity of plant virus derived DNA may be of less interest. Therefore, there is a need for easy, interactive, and in-depth browsing of the analysis results.

To that end the Virome Browser (Nieuwenhuijse, D. *et al*., 2019, in preparation) Shiny app was built, which allows users to download virus related analysis data form the PP and visualise and browse these locally. The Virome Browser allows users to import annotation data and assembled contigs from the PP, which can be interactively filtered based on quality thresholds such as amino acid identity and hit length. Subsequently, contigs with a specific annotation can be selected based on user interest. The selected contigs can then be inspected individually, allowing the user to visualise the sequence of the contig and the open reading frame (ORF) structure. Both the nucleotide and amino acid sequences derived from the ORFs can be saved as a FASTA file for further analysis. Metagenomic sequencing of the virome typically requires in-depth post-processing of the analysis results, which can be a daunting task for users without programmatic experience. The in depth analysis functionalities of the Virome Browser enable users without a bioinformatic background to extract useful information from raw analysis results. The Virome Browser has been made into an R package which can be downloaded and installed locally from github (currently private) using Rstudio.

## Configuration

As mentioned in the CDH design section, the Data Hubs allow for flexibility and choice of their configuration to cater for different collaborations. The configuration of a Data Hub depends on the data type being shared and analysed, and hence includes choices of an appropriate sample metadata standard as well as the analysis pipeline needed and the visualisation process. Both choices of sample metadata reporting standard (checklists, see also under “Upload and Standards”) and the analysis workflow depend on the sample type and provenance, for example whether the sequenced sample is a bacterial isolate, viral, or whether it is derived from an environmental setting.

## Usage and Access to the platform

Authenticated access via the PP interactive (https://www.ebi.ac.uk/ena/pathogens/) and the associated API (https://www.ebi.ac.uk/ena/portal/api/) enables the user to search pre-publication data in a desired Data Hub. To do this the user needs to authenticate using a username and password combination. The PP/API can also be used to access data in the public domain without user authentication. The search functionalities here are the same to when an user authenticates but results are limited to microbial data in the public domain. Figure III shows the use of the PP interactive interface going through steps of building a query to look in the public domain for read data of Ebola virus collected in Zaire in 1976, and customising the result page to return ‘centre name’, ‘study accession, ‘sample accession’, ‘run accession’ and ‘fastq_ftp’ (the FTP file path to the processed fastq files). The results can be downloaded in tab-separated value (TSV) and JSON formats.

**Figure III:**
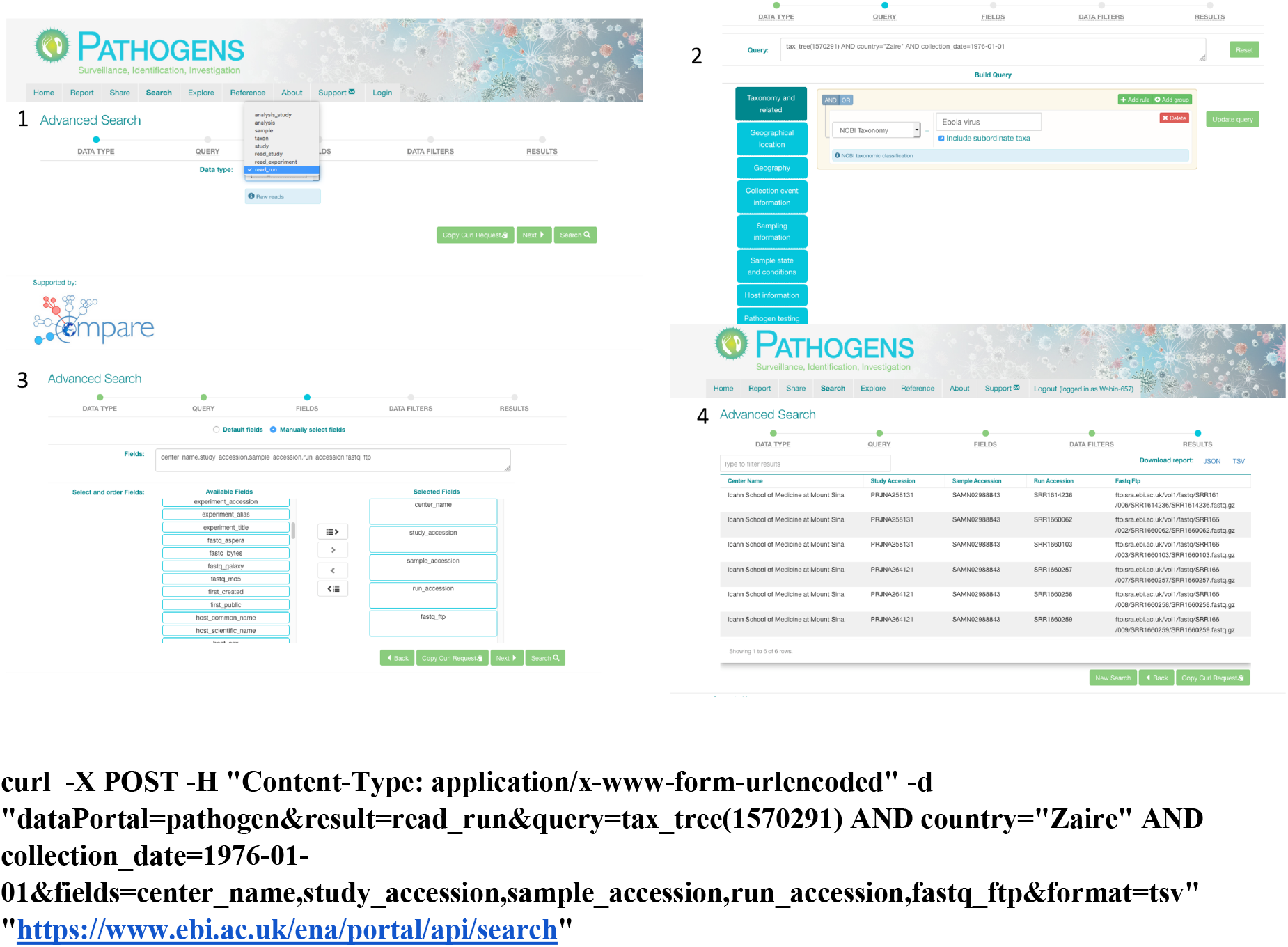
Pathogen Portal Interactive Interface. 1: select a domain (e.g. read_run); 2: narrow down search (where desired) by specifying taxonomy and sample collection details (e.g. collection date and country); 3: specify (where desired) which fields should be returned in the result report (e.g. center name, study accession, sample accession, etc); 4: result page with download options in TSV and JASON

In order to explore a data set that has been included in a Data Hub and run through one of the integrated analysis pipelines (SELECTA workflows) the user can go to the ‘Explore’ tab in the PP and visualise the analysis results via a connected Notebook. To demonstrate how this works, we selected three Salmonella enterica data sets that are available in the public domain. We configured a Data Hub, called dcc_benoit, to include this data and run it through the CGE bacterial analysis pipeline (BAP). The output of the analysis was used to create a public Notebook, the visualisation process has been rendered and can be accessed via the PP: https://www.ebi.ac.uk/ena/pathogens/explore and select the ‘View Demo’ button.

There are three tabs available here, ‘Data Hub content’ shows a summary tables for the data content of the Data Hub, ‘Primary Analysis’ shows in this case the visualisation of the CGE analysis pipeline output, and finally there is an ‘AMR’ tab here because antimicrobial resistance profiles (antibiograms) exist for 19 of the sequenced samples. For a demonstration video to see how to change views or filter data please go to https://www.ebi.ac.uk/ena/pathogens/explore and select the ‘View Demo’ button.

The COMPARE Data Hub model has been very successful and used for 13 different projects by the COMPARE collaboration. Use and assessment of the CDHs have been also described in Poen M. *et al.* (2019), and Matamoros S. *et al.* (2019), both submitted. Table II lists CDHs and their descriptions. Where data is already in the public domain it is indicated by the corresponding URL to ENA. In other cases data is still pre-publication confidential at the time of writing.

**Table II:** COMPARE Data Hubs

| Name | Topic | Description |
| --- | --- | --- |
| dcc_sibelius | Pilot: Influenza H5N8 | This datahub contains RNA sequence data and metadata of three highly pathogenic avian influenza H5N8 viruses used in the H5N8 pilot project. The aim of this project is to determine the similarity and reproducibility of viral genome consensus determination and minority variant detections between three different but commonly used workflows.<br>*Data public and described in Poen M., et al. (2019), submitted.* |
| dcc_berlioz | Pilot: Ebola | This datahub contains Ebola sequence data generated as part of a relief effort in West Africa. Data public at: https://www.ebi.ac.uk/ena/data/view/PRJEB10265 |
| dcc_liszt | Global Sewage | This datahub contains the DNA sequences, metagenomics data from the Global Sewage Surveillance project. The data represent all urban sewage samples collected prior to treatment plant inlet from major cities around the world. The study aims to establish a global surveillance of infectious disease agents and AMR. |
| dcc_strauss | Kibera Sewage | This datahub contains the DNA sequences, metagenomics data from the informal settlement of Kibera, Nairobi, Kenya. The data represent sewage samples collected over 3 months from two of the 10 population clusters under US CDC surveillance. The study aims to establish a hot spot disease surveillance. |
| dcc_vivaldi | Foodborne pathogen surveillance and epidemiological analysis | This datahub contains the DNA sequences and metagenomics data from the COMPARE Work package 4/7 workgroup of food-borne pathogens. The majority of the data consist of the bacterial pathogens Salmonella, E coli, Listeria and Campylobacter. Samples are collected from human infections, food and animals and the work done on these sequences aimed to improve and evaluate new and established microbiological and epidemiological analysis methods used in Public Health and the food and veterinary sector. Data were also collected with the goal of developing Source Attribution methods for WGS data. |
| dcc_schubert | AMR working group | This datahub contains DNA sequences of bacterial isolates' genomes coupled with phenotypic antimicrobial resistance (AMR) data. The phenotypic data are presented in standardized antibiograms, describing for each isolate the antibiotics tested, the levels of resistance observed, the antimicrobial susceptibility testing method employed, etc. The aim is to provide a platform for exchange of genomic and phenotypic information regarding AMR, thus encouraging surveillance and the development of innovating projects for the prediction of AMR from sequence data.<br>*Data public and described in Matamoros S., et al. (2019), submitted.* |
| dcc_brahms | Diagnostic metagenomics on clinical samples | Diagnostic metagenomics on clinical samples: Prediction of antibiotic resistance genes and pathogen discovery in shot-gun metagenomic data from swine feces.<br>The data-hub contains shot-gun metagenomic NGS data sets generated from swine holdings with acute diarrhea. The causative agent, Porcine epidemic diarrhea virus (PEDV), was identified. Metadiagnostic analysis via FLI RIEMS pipeline revealed co-infection with bacteria. Prediction of antibiotic resistance markers and pathogen discovery will be done as a pilot to test pipelines and algorithms with clinical metagenomics data. |
| dcc_handel | Virus metagenomics | This datahub is used to share Fastq files of NGS experiments, mostly with a metagenomics approach, on clinical samples of patients with hepatitis A and norovirus gastroenteritis. The human reads in the files have been removed before being uploaded. |
| dcc_puccini | Parasites (Comparative Genomics of Intestinal Protozoa) | This datahub contains the raw DNA sequences from isolates of the protozoan *Cryptosporidium*. The data were generated in collaboration with the UK *Cryptosporidium* Reference Unit, and represent both sporadic and outbreak cases. The study aims to understand the major factors that structure parasites' genomes by using a comparative genomics approach.<br>Data public at:<br>https://www.ebi.ac.uk/ena/data/view/PRJEB15112 |
| dcc_beard | Global Sewage snapshot Virome sequencing part | This datahub contains the raw read data of the virus specific part of the Global Sewage Surveillance project. The data aim to capture the complete DNA and RNA virome of the sampled locations. In addition, the analysis results of the SELECTA-SLIM pipeline can be found in this datahub. |
| dcc_cole | Metagenomics ring trial | This datahub contains the DNA sequences, metagenomics data from the Food Metagenomics ring trial 2018. The data represent DNA- and RNA-derived metagenomics data sets processed from a piece of smoked salmon spiked with a complex mock community consisting of viruses, bacteria, fungi and a parasite. The study aims to compare wet lab protocols using the same starting material. |
| dcc_bromhead | CoVetLab (Colistin resistant Enterobacteriaceae project) | This datahub contains the DNA sequences of single bacterial isolates of primarily antimicrobial resistant enterobacteriaceae from European National Reference Laboratories. The data represent among others isolates collected for the EU antimicrobial resistance monitoring in zoonotic and indicator bacteria from humans, animals and food as well as for the CoVetLab, colistin resistance project. The intention is that all data related to the EURL-AR will be submitted to the data hub. |
| dcc_schumann | Bioaccumulation experiment | This datahub contains DNA sequences and metadata to analyze bioaccumulated oysters. In addition, the analysis results of the SELECTA-SLIM pipeline can be found in this datahub. |

## Data Hub Requests

Requests to configure a Data Hub with a short description of the project/collaboration should be sent to. If requesting groups know already which scope they wish to use, i.e. sample metadata standard and analysis pipelines described in this work, these should be also specified. Alternatively, design of new metadata standards and integration of other analysis workflows will become part of a consultation process. We will send out a document to be completed by the requesting group to include names, affiliations and contact details of data providers and users who will need to provide consent to access pre-publication data (if required). Once the scope of the Data Hub is finalised a Data Hub will be assigned and configured accordingly. Following this process users will be sent access credentials to their assigned Data Hub. Data providers Webin accounts will be associated with the Data Hub, so they can use the PP to share submitted data with other Data Hub users (see also under ‘Sharing’).

## Data Hub Sharing Agreements

Considering the governance of access and use of data within the COMPARE Data Hubs, and in order to address concerns of the Parties involved in terms of pre-publication access, confidentiality, due diligence (compliance with relevant regulations), and clarifying specific rights and obligations, a Code of Conduct was agreed by the COMPARE members in the Consortium Agreement. Additionally, in order to set up pilot projects that were relevant to the COMPARE platform and required the use of Data Hubs but were executed by external users (non Consortium members), the Code of Conduct (http://www.compare-europe.eu/project-organisation/work-packages/workpackage-12) had to be signed by all parties involved in order to gain access and permission to use the available data for the purpose of specified activities. Within this Code of Conduct, the confidentiality and due diligence agreement is stating the ownership of, and responsibilities for the sharing of data in the Data Hubs, in addition to the Terms of Reference that clarify the do’s and don’ts for the participating parties. The ultimate goal of this documents is to protect the integrity of the developed database and the Platform as a whole, to promote open access but facilitate also temporally confidential sharing when needed, and promote constructive peer collaboration.

Efforts have been made to draw up the agreements in a way that is legally unambiguous and at the same time readable and understandable for participants who are not legally qualified. Another special consideration was made on the flexibility of terms, leaving to the participants to decide, at any stage, on the issues of a) who can participate: parties can join (if agreed by all participants) and/or drop off at any point in time; and b) the nature of data to be shared: the amount and type of metadata attached to the raw sequences, considering a defined minimum set of metadata (see also under “Upload and Standards”). The main concerns of the stakeholders involved addressed on the Code of Conduct are stratified between the two defined phases: 1) protected space, to which only eligible participants who have undersigned the agreements have access, and 2) public availability of the data. For the first stage, issues of confidentiality and ownership are directly addressed, when Parties agree not to share the data outside the closed space, and not use the data in commercial applications and/or scientific publications without the consent and acknowledgement of the data providers. For the second stage, to comply with open data policies and as a preemptive response to requests of third parties to the publication of data, Parties declare to share the data on the public domain with a minimum set of metadata. At both stages, Parties declare to refrain from any attempt to identify individuals when using the data, and that data is uploaded and available in accordance with the applicable laws and regulations of the European Union and of the country of origin.

## Ongoing developments

The CDHs have been implemented over the last several years in an approach that saw a simple first implementation with rapid new deployments as additional functions became available. We will continue with this approach in order to provide maximum benefit as early as possible to our users. Particular areas of focus over the next year will likely be updates to analysis workflows, tools to search based on queries within the outputs of analyses, such as species identified, typing information and resistance gene calls, searchability within AMR resistance profile antibiograms and implementation of the Evergreen tree-building system (https://cge.cbs.dtu.dk/services/Evergreen/, Szarvas J. *et al.* submitted, pre-print: https://www.biorxiv.org/content/10.1101/540138v1).

A current priority is to pilot the system in a variety of contexts, spanning pathogen, sectors and domains, including public and animal health agencies, food safety monitoring organisations and the commercial food industry. We invite interested groups from these and other contexts to contact us with ideas for pilots.

## Author contributions

Clara Amid led on the development of the manuscript; Guy Cochrane, Marion Koopmans and Frank Møller Aarestrup conceived the CDH; Clara Amid, Sam Holt, Jeffrey E. Skiby operated user support systems; Nima Pakseresht, Blaise Alako and Nadim Rahman provided cloud compute environment used in CDH; Nicole Silvester, Peter Harrison, Suran Jayathilaka and Abdulrahman Hussein provided metadata search, data access APIs and access management tools; Ole Lund and Lukasz D. Dynovski provided the DTU Uploader; Rasko Leinonen provided the Webin data submission system; Ole Lund, Jose J. L. Cisneros, Rolf S. Kaas, Martin C. F. Thomsen and Camilla Hundahl provided the CGE analysis workflow; Dirk Höper and Ariane Belka contributed the RIEMS analysis workflow; István Csabai, Dávid Visontai, Bálint Á. Pataki, József Stéger and János M. Szalai-Gindl supported data visualisation; Matthew Cotten and David Nieuwenhuijse contributed the SLIM analysis workflow and visualisation system; Surbhi Malhotra-Kumar and Basil Britto Xavier provided the BacPIPE analysis workflow; and George B. Haringhuizen and Carolina dos S Ribeiro provided user data agreement support.

